# A paired analysis of mercury among non-invasive tissues to inform bat conservation monitoring

**DOI:** 10.1101/2024.03.31.587502

**Authors:** Molly C Simonis, Kimberlee Whitmore, Kristin E Dyer, Meagan Allira, Bret Demory, Matthew M Chumchal, Daniel J Becker

## Abstract

Contaminant exposure can harm wildlife. However, measuring contaminant exposure in wildlife can be challenging due to accessibility of species and/or sampling tissue matrices needed to answer research questions regarding exposure. For example, in bats and other taxa that roost, it may be best to collect pooled feces from colonies for minimal disturbance to species of conservation concern, but fecal contaminant concentrations do not provide contaminant bioaccumulation estimates. Thus, there is a need for quantifying relationships between sample matrices for measuring contaminant exposure to answer research questions pertaining to wildlife health and addressing conservation needs. Our goal was to determine relationships between fecal and fur total mercury (THg). To do so, we collected paired feces and fur from Mexican free-tailed bats (*Tadarida brasiliensis*) in summer 2023 in western Oklahoma at a maternity roost with no known Hg point source. We analyzed THg in each sample matrix for each individual (n = 48). We found no relationship between individual fecal and fur THg. However, when averaged, fur THg was 6.11 times greater than fecal THg. This factor can be used as a screening-level risk assessment of under-roost feces, which could then be followed by direct assessments of fur THg concentrations and health impacts. We encourage the use of this conversion factor across other insectivorous bat species and sites for estimating initial risks of contaminant exposure with minimal disturbance to species of conservation concern, when timely research for conservation actions are needed, and when a contaminant point source is not yet known.

**Graphical Abstract:** 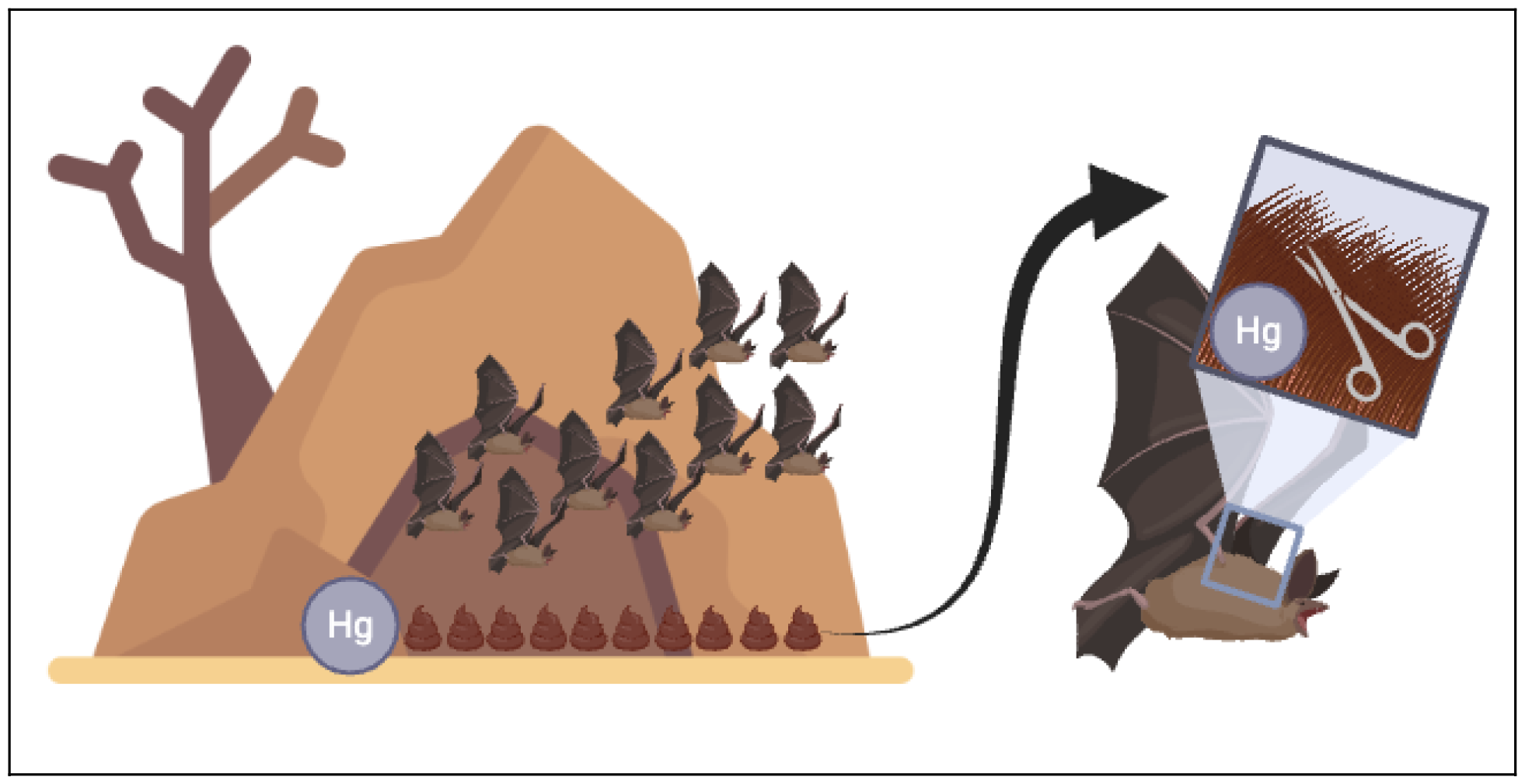

Graphical abstract created in BioRender under a free subscription. Cave icon created by artist Freepik at https://www.flaticon.com/free-icons/cave.

**Highlights:**

- Under-roost sampling for contaminant exposure minimizes species disturbance
- Contaminant exposure relationships in tissues can aide in measuring wildlife health
- We sampled *Tadarida brasilliensis* for paired fecal and fur total Hg (THg)
- THg in fur averaged 6.11 times greater than feces
- This factor can be used as an initial risk assessment for under-roost fecal sampling
- More invasive follow-up sampling (bat fur) can be justified following risk assessment

## 1. Introduction

Exposure to high concentrations of heavy metal contaminants such as mercury (Hg) can harm wildlife (Chételat et al., 2020). For example, at elevated concentrations, Hg exposure can reduce reproductive success, cause immunotoxicity, and damage genetic material (Wolfe et al., 1998). To determine how contaminant exposure affects wildlife health, researchers must be able to sample the appropriate tissue matrix (e.g., fur, blood, tissue, feces) that best relates to the health outcome of interest, as different matrices indicate different contaminant exposure and/or bioaccumulation timelines (Chételat et al., 2020). For example, Hg concentrations in fur represent longer-term exposure (e.g., weeks to months depending on molting cycles), while Hg concentrations in feces represent more acute exposure/ingestion of Hg (e.g., days to weeks depending on metabolism and excretion efficiency; Chételat et al., 2020). Choosing a matrix for contaminant sampling of wildlife may also be complicated by instances where direct sampling is not possible, such as wanting to minimize disturbance when sampling species of conservation concern. Therefore, there may be a mismatch between the sampling matrix needed and the sampling matrix available for measuring wildlife health.There remains a need to understand how contaminant concentrations across sampling matrices relate to one another to optimize monitoring practices that inform wildlife health and conservation research.

Bats are a highly diverse group of mammals that are of special interest to conservation (Kunz and Fenton, 2003). Bats are important bioindicator species due to their long lifespans (e.g., the oldest known wild bat was 41 years old; Podlutsky et al., 2005), slow birth rates (one to two pups per year; Wilkinson and South, 2002), high mobility through flight, diverse habitat use, and—for insectivorous bats—feeding at high trophic levels (Jones et al., 2009; Russo et al., 2021; Zukal et al., 2015). In North America specifically, many bats are threatened by emerging infectious diseases, namely white-nose syndrome (Blehert et al., 2009; Lorch et al., 2011). Exposure to contaminants such as Hg can increase bat infection risk (Becker et al., 2021) and can shape population-level infection prevalence through diverse feedback loops (Becker et al., 2024; Cable et al., 2021).

Depending on the bat species, collecting samples needed to quantify health can be complicated (Giles et al., 2021), particularly when interested in contaminant concentrations. For example, sampling pooled bat feces or urine under roosts or outside cave entrances may be most appropriate for species of conservation concern to minimize disturbance (Giles et al., 2021; Walker et al., 2016); however, feces can only provide insight into short-term contaminant exposure, and pooled sampling is not traceable to individuals. Estimating long-term contaminant exposure is better measured by sampling bat fur, which represents bioaccumulation in insectivorous bats due to its strong correlation with contaminants in blood (Wada et al., 2010; Yates et al., 2014). In cases where sampling fur from bats is logistically challenging (e.g. disturbance of threatened and endangered species), researchers could collect under-roost fecal samples as a preliminary screening assessment before moving toward increasing disturbance for fur collections. However, formal methods for converting fecal contaminant concentrations to bioaccumulative measures in fur are lacking, hindering our ability to identify bat health risks in a timely manner and, in turn, delaying conservation actions.

Here, we aim to 1) assess Hg in bat feces and fur and determine relationships between the two concentrations, representing the potential to estimate long-term Hg exposure from acute Hg exposure; and 2) determine if effects of body mass on bioaccumulation are detectable in feces, as expected for fur (Mina et al., 2019; Timofieieva et al., 2021; Yates et al., 2014). We expected that overall Hg concentration in fur would be greater than those in feces across paired samples from individual insectivorous bats, and that fur Hg concentrations would increase with fecal Hg concentrations. We expected these differences due to bioaccumulation timelines associated with repetitive tissue turnover (fur) vs. excretion (feces). We also expected fecal and fur Hg concentrations to increase with body mass, representing bioaccumulation of Hg from long-term exposure (fur) and large bats ingesting more Hg-exposed insects (feces). Results presented here can inform whether fecal Hg can serve as a substitute for fur Hg, such as in cases where pooled under-roost sampling is the primary option for quantifying contaminants in bats. Thus, our results have the potential to inform monitoring strategies and design of research questions regarding contaminant exposure in bats.

## 2. Materials and Methods

### 2.1 Study site and species

As part of a larger study of bat migration, immunity, and infection (Becker et al., 2024), we quantified Hg concentrations in Mexican free-tailed bats (*Tadarida brasiliensis*) at the Selman Bat Cave in Woodward County, Freedom, Oklahoma, USA. *T. brasiliensis*, a common insectivorous bat across North America, migrates long-distances annually in spring and fall between northern and southern portions of their range (Wilkins, 1989). These bats are highly valuable to conservation due to the ecosystem services they provide throughout their range. *T. brasiliensis* provide important nutrient transfer to soils across the USA and Mexico from their guano, particularly in the USA where they form large maternity colonies in spring and summer (thousands to millions of individuals; Bernardo and Cockrum, 1962). In the USA, *T. brasiliensis* eat up to 79% of their body weight in agricultural pests (about 8.1 g of insects per individual) each night during peak lactation (Kunz et al., 1995), saving up to $1.7 million in agricultural pest control each year (Cleveland et al., 2006). Finally, *T. brasiliensis* are valuable to the ecotourism industry, with public viewing of nightly maternity colony emergence throughout the southwestern USA estimated to bring in over $6.5 million to local economies (Bagstad and Wiederholt, 2013), highlighting their high intrinsic and monetary value to the public.

### 2.2 Bat capture

At the Selman Bat Cave, we captured 48 *T. brasiliensis* on 16 June, 17 June, and 18 July 2023. *T. brasiliensis* were captured with hand nets upon emergence from the cave and temporarily held in individual cloth bags. Most *T. brasiliensis* arrive at Selman Bat Cave in April each year following long-distance migration from Mexico (Glass, 1982). Capturing bats in June and July ensured 1) bats were foraging within the local landscape for multiple months and 2) fur had grown during their summer residency (Constantine, 1957; Fraser et al., 2013). From each bat, we collected paired samples of 1-2 fecal pellets (opportunistically collected from bat holding bags) and fur trimmed from the mid-dorsal region. We weighed each individual to the nearest gram and identified bats by sex, reproductive status (i.e., non-reproductive, testes descended, pregnant, lactating, and post-lactating), and age (i.e., juvenile, subadult, and adult; Morgan et al., 2019). Sampling was approved by the Institutional Animal Care and Use Committee of the University of Oklahoma (2022-0198), with a collection permit from the Oklahoma Department of Wildlife Conservation (10567389). All bats were released after sampling. Feces per individual were stored separately in 1 mL 95% ethanol in a portable -20 °C cooler, and individual fur samples were stored in 1.5 mL microcentrifuge tubes at room temperature. Samples were transported to the University of Oklahoma, where feces were stored at -20 °C and fur remained at room temperature.

### 2.3 Total mercury analyses

At the University of Oklahoma, we prepared individual fur samples to perform total Hg (THg) analyses (MeHg and inorganic Hg combined). The mass of fur used ranged from 0.8–4.9 mg, with over 75% of samples having at least 4 mg fur (11 samples had smaller volumes). Once weighed, fur samples were heat treated at 60 °C for 30 min. Feces (on dry ice) and fur (at room temperature) were then shipped to Texas Christian University for THg analyses.

At Texas Christian University, all fecal and fur samples were analyzed for THg using direct Hg analysis (Nippon MA-3000, NIC), which uses thermal decomposition, gold amalgamation, and atomic absorption spectroscopy (US Environmental Protection Agency, 2018). Quality assurance included reference (National Research Council of Canada Institute for National Measurement Standards) and duplicate samples. Reference samples (TORT-3 and PACS-2) were analyzed every 20 samples, and the mean recovery percentage for TORT-3 was 99.1 ± 2.42 % (n = 13). The mean recovery percentage for PACS-2 was 98.9 ± 5.74 % (n = 13). Duplicate samples of fur were analyzed every 20 samples, and the mean relative percent difference was 1.86 ± 1.78 % (n = 6). Duplicate samples of feces were analyzed every 20 samples, and the mean relative percent difference was 7.59 ± 3.24 % (n = 2). The limit of detection was determined by utilizing the measured limit of blank and test replicates of a sample known to contain a low concentration of analyte (Armbruster and Pry, 2008). All samples were above the limit of detection of 0.02 ng THg.

### 2.4 Statistical analyses

We performed all statistical analyses and data visualization in the statistical environment R version 4.3.1 ‘Beagle Scouts’ (R Core Team, 2023). We created a generalized linear mixed effects model (GLMM) with a Gamma family and log link using the function glmer() from the *lme4* package (Bates et al., 2015) to determine differences between paired bat fecal and fur Hg concentrations. This GLMM was fit to THg concentrations (mg/kg) as a function of sample matrix (feces or fur) and random effects of the individual bat and date of sample collection (models with nested random effects did not converge). We tested differences between paired samples using a Type II ANOVA with the function Anova() from the *car* package (Fox and Weisberg, 2019), and obtained estimated marginal means of fur and fecal THg using the package *emmeans* and its self-titled function (Lenth, 2021). We also obtained marginal and conditional R^2^ for this GLMM using the function r.squaredGLMM() from the package *MuMIn* (Bartoń, 2023). To determine if there was a direct relationship among individual bats between THg in feces and fur, we fit another GLMM with a Gamma family and log link for THg in fur as a function of THg in feces with a random effect of date of sample collection, using the glmer() function in *lme4* (Bates et al., 2015). We tested main effects with a Type II ANOVA and assessed model fit using functions Anova() and r.squaredGLMM() in the *car* and *MuMIn* packages (Bartoń, 2023; Fox and Weisberg, 2019). Finally, to determine if THg in feces and/or fur provided evidence of Hg bioaccumulation, we fit another GLMM with a Gamma family for THg as a function of an interaction between mass (g) and sample matrix (feces or fur) and random effects of individual bat and date of sample collection (Bates et al., 2015), again testing main effects with a Type II ANOVA with the Anova() function via the *car* package (Fox and Weisberg, 2019), and quantifying adjusted R^2^ using the r.squaredGLMM() function from the package *MuMIn* (Bartoń, 2023). All data visualizations were created using the package *ggplot2* (Wickham, 2016).

## 3. Results

We captured 48 *T. brasiliensis* representing 44 females (five non-reproductive, nine pregnant, 25 lactating, five post-lactating), three males (all non-reproductive juveniles), and one bat whose sex was unknown. Of the captured bats that were female, two were juveniles, one was a subadult, and 41 were adults. We captured most bats for paired feces and fur samples during June dates (18 June 2023 n = 22; 19 June 2024 n = 12), with the remaining bats sampled on 18 July 2023 (n = 14). Regardless of sample matrix, average THg was less than 1 mg/kg (mean ± SE; 0.66 ± 0.09).

Average fecal THg was 0.19 ± 0.01 mg/kg (raw mean ± SE) and average fur THg was 1.14 ± 0.07 mg/kg. When comparing THg across pooled sample matrices, our GLMM indicated significant differences between sample matrices (χ^2^_1,47_ = 764.12, P < 0.001, R^2^_m_ = 0.83, R^2^_c_ = 0.88; Figure 1), such that concentrations in bat fur were more than six times greater than feces (mean [lower CI, upper CI]; feces: 0.18 [0.16, 0.21] mg/kg, fur: 1.087 [0.95, 1.25] mg/kg). We found no direct relationship between fecal and fur THg across individuals (χ^2^= 0.26, P = 0.62, R^2^_m_ = 0.01, R^2^_c_ = 0.14; Figure 2). Finally, the relationship between bat mass and THg did not differ by sample matrix (χ^2^ = 0.00, P = 1.00, R^2^_m_ < 0.01, R^2^_c_ =1.00; Figure 3), with random R^2^_c_ effects of individual bat and sample collection date accounting for almost all variation within the model (as indicated by R^2^_c_). Additionally, neither sample matrix nor mass influenced THg independently (sample matrix: χ^2^ = 0.00, P = 1.00; mass: χ^2^ = 0.00, P = 1.00; Figure 3).

**Figure 1.**
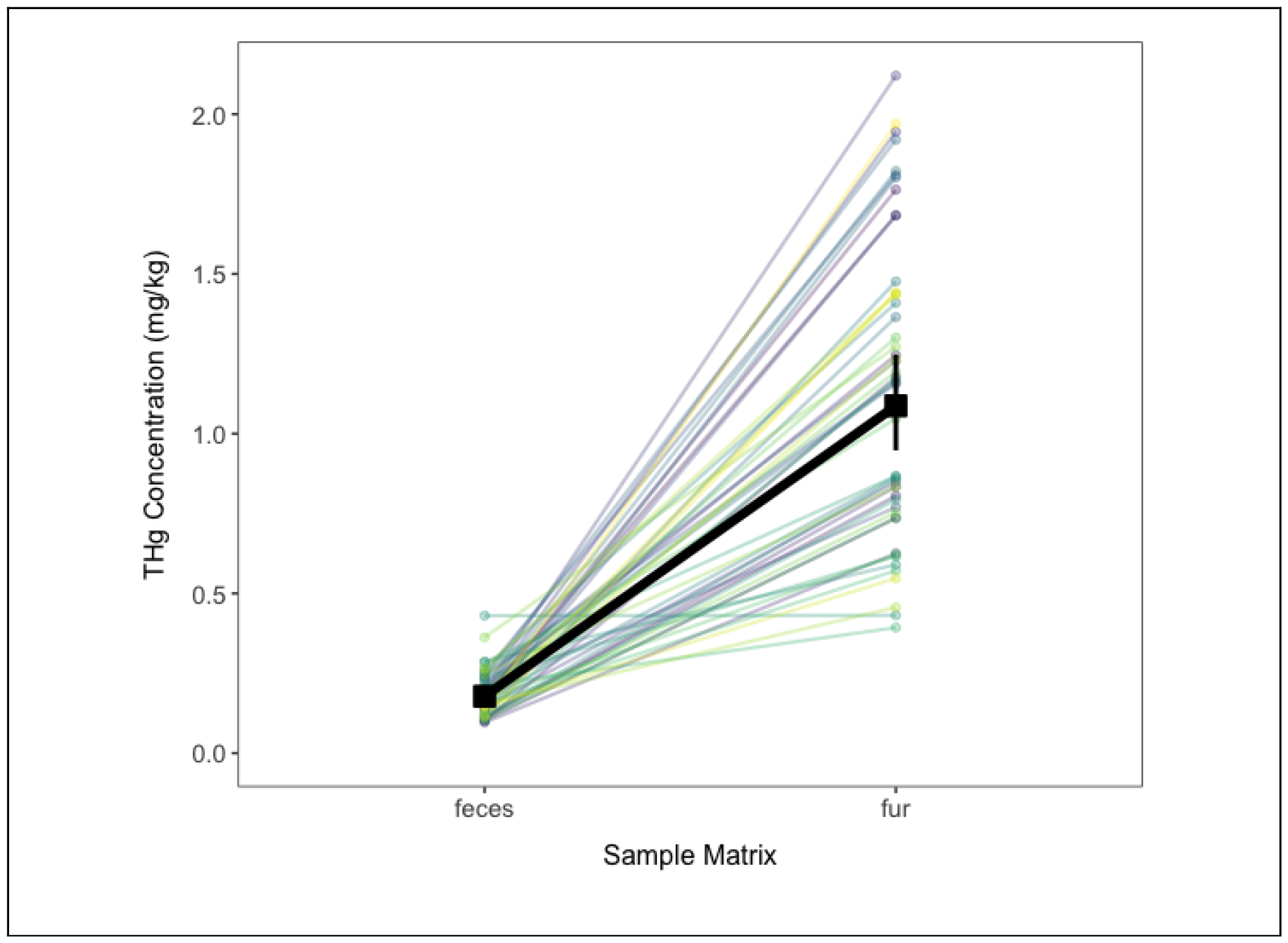
THg concentrations in *T. brasiliensis* fur were 6.11 times greater than those in feces. Fecal and fur samples were paired with each individual, as represented by lighter colored points and lines. Estimated marginal means and 95% confidence intervals of THg concentrations are represented in thicker black points and corresponding lines.

**Figure 2.**
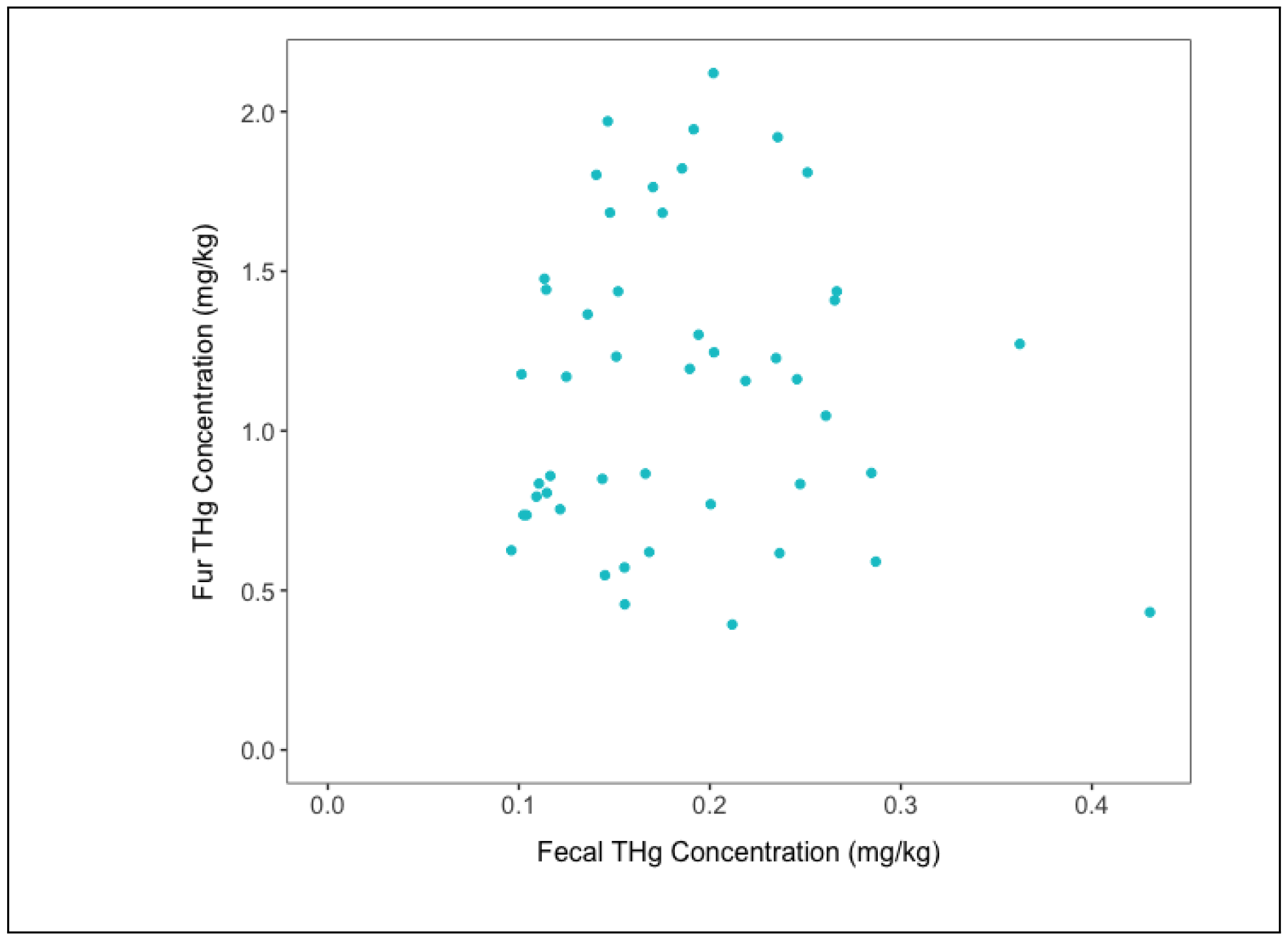
There was no relationship between fecal and fur THg across individual *T. brasiliensis*. Fecal and fur samples were paired with each individual, as indicated by individual points.

**Figure 3.**
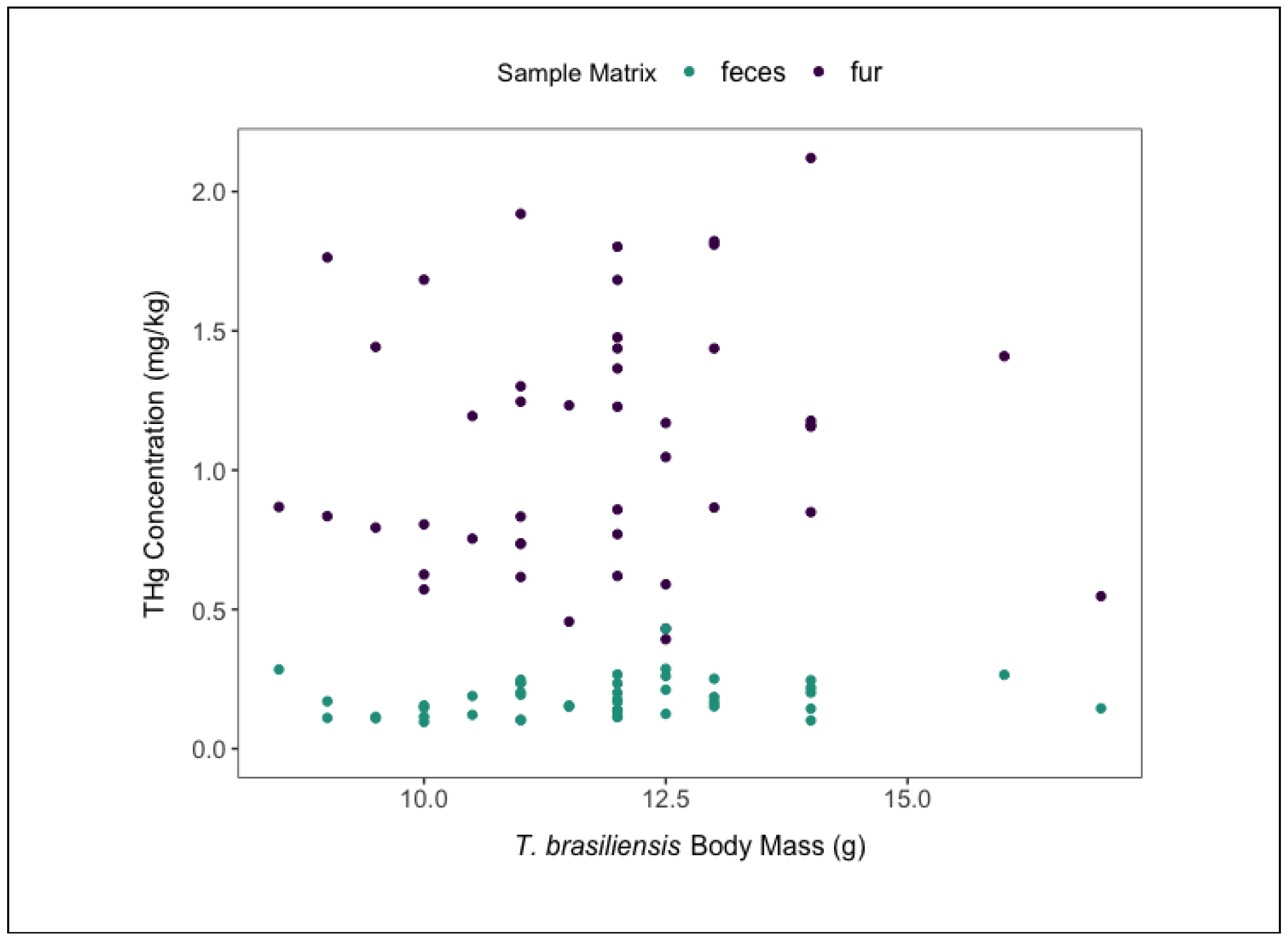
We observed no relationship between *T. brasiliensis* body mass and THg concentrations by sample matrix, across or within each sample matrix. Colors indicate different sample matrices measured for paired sampling of each individual bat.

## 4. Discussion

Determining relationships between short- and long-term contaminant exposure can benefit wildlife research and conservation, as indirect samples representing short-term exposure such as feces may often be more readily accessible than long-term exposure samples requiring direct capture of threatened species, such as fur. Using paired samples from individual *T. brasiliensis* in the Great Plains of the United States, we found THg concentrations in bat fur were on average 6.11 times greater than THg concentrations in feces. However, we found no relationship between individual fecal and fur THg concentrations, nor did body mass explain variation in THg within or across sampling matrices. Taken together, these results suggest that feces should not be used as a substitute for sampling other matrices such as fur to determine long-term contaminant risks in bats. However, fecal THg concentrations could be used to estimate fur THg concentrations with our estimated conversion factor of 6.11 for initial risk-level assessments to determine if further invasive sampling is needed, such as if estimated fur THg concentrations are above acknowledged thresholds for negative health effects..

Fecal and fur THg concentrations measured here are similar to concentrations found in other *T. brasiliensis* populations and other insectivorous bat species in North America. In the northeastern US, nine species of insectivorous bats had THg concentrations in fur below 1 mg/kg (Yates et al., 2014). *T. brasiliensis* across two sites in Texas averaged fur THg concentrations below 2 mg/kg (Korstian et al., 2018). Pooled feces collected under bat boxes mostly occupied by *T. brasiliensis* in Florida also had average THg concentrations below 1 mg/kg (Edwards et al., 2019). When averaged, our THg concentrations from *T. brasiliensis* feces and fur were 0.19 ± 0.01 mg/kg and 1.14 ± 0.07 mg/kg, respectively. Fur THg values here are also comparable to fur THg concentrations in *Eptesicus fuscus* (big brown bats) captured at reference sites in the eastern USA (Wada et al., 2010). Similar to these reference values, we are also not aware of any immediate point source of Hg, such that THg values in Great Plains *T. brasiliensis* are likely reflective of atmospheric deposition (Chételat et al., 2018; Korstian et al., 2018). Taken together, our results suggest that a conversion factor of 6.11 could be used to estimate fur THg concentrations from fecal THg concentrations, which may be most applicable for bat species of conservation risk where under-roost sampling is preferable to direct capture to minimize disturbance. However, given the lack of an individual-level relationship between fecal and fur THg concentrations, we suggest this conversion factor should only be used to assess whether fur THg could be at concentrations indicative of health risks, and thus, warranting further invasive sampling (e.g. fur, blood) for confirmation of contaminant concentrations and assessment of conservation and health risks.

In mammals, lethal effects of Hg toxicity typically occur above concentrations of 10 mg/kg in tissues such as brain, liver, kidney, and fur (Nam et al., 2012; Shore et al., 2011; Wolfe et al., 1998). Thus, a threshold of estimated fur THg around 10 mg/kg following pooled fecal sampling and conversion by a factor of 6.11, would indicate an immediate need for fur samples and conservation action. However, non-lethal thresholds of Hg exposure below 10 mg/kg are well-reported in mammals and bats specifically, but still with potentially adverse health effects. For example, Neotropical bat cellular immunity and bacterial killing ability are negatively correlated with THg concentrations below 10 mg/kg (Becker et al., 2021, 2017). Additionally, in fruit-eating Neotropical bats, the probability of infection with bartonellae (common bacterial pathogens) was associated with higher fur THg, even when concentrations were as low as 0.3 mg/kg (Becker et al., 2021). Neotropical bats with the highest spleen (0.25 mg/kg) and liver (0.23 mg/kg) THg concentrations also had the highest micronuclei intensities in red blood cells (Calao-Ramos et al., 2021), representing genotoxic effects of environmental pollutants (Samanta and Dey, 2012; Sandoval-Herrera et al., 2021). All of our THg concentrations were below 2.2 mg/kg (max THg = 2.1 mg/kg in an individual fur sample). While our THg concentrations in *T. brasiliensis* feces and fur are relatively low, and no known Hg point source exists near our sampling site, there could still be concern for adverse health effects associated with Hg exposure. Wildlife managers would need to make decisions regarding a potential THg thresholds for initial risk assessment with pooled fecal sampling to determine when more invasive fur sampling outweighs disturbance risk, particularly for species of conservation concern. To help determine non-lethal thresholds of Hg exposure that may have adverse health effects, future research priorities should include assessing immunity, infection, and genotoxic correlates of Hg exposure in this and other insectivorous bat species important to wildlife conservation.

We did not find relationships between THg and body mass in either sampling matrix, even though we expected fur THg to be positively correlated with body mass, due to its better indication of bioaccumulation (Yates et al., 2014). As noted above, we also did not detect an individual relationship between fecal and fur THg, which could be due to the greater variability in fur THg. First, our sampled *T. brasiliensis* included adult (n = 41), subadult (n = 1), and juvenile bats (n = 5). While sample sizes for subadults and juveniles did not allow for statistical comparisons, different age groups could have introduced greater variation in fur THg (Yates et al., 2014). Second, juvenile fur growth occurs in late summer in the maternity colony, while there may be more variability in adult and subadult molt and fur growth timing (Fraser et al., 2013). We sampled *T. brasiliensis* during times that both adults and juveniles should have fur growth and thus, should represent bioaccumulation. However, fur THg from the July sampling date was relatively lower than both June sampling dates, suggesting that some individuals may have molted later in the summer than expected and indicating losses of cumulative THg across progressive sampling periods. Although THg concentrations can vary seasonally (Albert et al., 2021; Yates et al., 2014), we accounted for sample collection date variation by including it as a random effect in our GLMMs. Yet sample collection date accounted for low proportions of residual variance within our GLMMs (i.e. 6% for the GLMM in Figure 1, 13% for the GLMM in Figure 2, and <1% for the GLMM in Figure 3). Increasing paired sampling in the future could detect expected increases of fur THg with body mass and better identify differential relationships between age classes and molt timing across the summer maternity season.

## 5. Conclusion

In conclusion, understanding relationships between contaminant concentrations across sampling matrices can inform diverse applications in wildlife conservation research and management. Our study estimated a conversion factor between fecal and fur THg concentrations in a common insectivorous bat species, enabling researchers to extrapolate fur concentrations when only fecal samples are logistically feasible for initial risk assessments. We encourage the use of this conversion factor across other insectivorous bat species and sites when species of conservation concern require initial screening-level risk assessment sampling with minimal disturbance. Future research should include identifying immune and infection correlates of fecal and fur THg concentrations to identify the health implications of contaminant exposure and evaluate needs for conservation action.

## Acknowledgements

This work was supported by the National Institute of General Medical Sciences of the National Institutes of Health (P20GM134973) and Research Corporation for Science Advancement (RCSA). This work was conducted as part of Subaward No. 28365, part of a USDA Non-Assistance Cooperative Agreement with RCSA Federal Award No. 58-3022-0-005. MCS was supported by an appointment to the Intelligence Community Postdoctoral Research Fellowship Program at University of Oklahoma administered by Oak Ridge Institute for Science and Education through an interagency agreement between the U.S. Department of Energy and the Office of the Director of National Intelligence. We thank the Oklahoma Department of Wildlife Conservation for site access and the Selman Living Laboratory for fieldwork support. We also thank Lauren Lock, Jaleel Zubayr, and Drake Johnston for assistance in the field and preparing samples for THg analysis.

## Data Availability

All data and R code used for analyses are available on MCS’ Github page (username: simonimc; https://github.com/simonimc/Tadarida_brasiliensis_paired_fecal_fur_THg).

